# Prism adaptation modulates connectivity of the intraparietal sulcus with multiple brain networks

**DOI:** 10.1101/683078

**Authors:** Selene Schintu, Michael Freedberg, Stephen J. Gotts, Catherine A. Cunningham, Zaynah M. Alam, Sarah Shomstein, Eric M. Wassermann

## Abstract

Prism adaptation (PA) alters spatial cognition according to the direction of visual displacement by temporarily modifying sensorimotor mapping. Right-shifting prisms (right PA) improve neglect of left space in patients, possibly by decreasing activity in the left hemisphere and increasing it in the right. Left PA shifts attention to the right in healthy individuals by an opposite mechanism. However, functional imaging studies of PA are inconsistent, perhaps because of differing activation tasks. We measured resting-state functional connectivity (RSFC) in healthy individuals before and after PA. Right, vs. left, PA decreased RSFC in the navigation network defined by the right posterior parietal cortices (PPCs), hippocampus, and cerebellum. Right PA, relative to baseline, increased RSFC between regions within both PPCs and between the PPCs and the right middle frontal gyrus, whereas left PA decreased RSFC between these regions. These results show that right PA modulates connectivity within the right-hemisphere navigation network and shifts attention leftward by increasing connectivity in the right frontoparietal network and left PA produces essentially opposite effects, consistent with the interhemispheric competition model. These finding explain the action of PA on intact cognition and will help optimize interventions in neglect patients.

---

Adaptation to prisms, which shift vision laterally, temporarily modifies sensorimotor mapping (Helmholtz, 1867; Rossetti et al., 1998). Prism adaptation (PA) is performed by practicing pointing movements to a displaced target, resulting in a rapid correction of the pointing error and a corresponding aftereffect in the opposite direction after the prisms are removed. Those aftereffects are not limited to the sensorimotor domain, but also affect cognition (Rossetti et al., 1998).

Adaptation to right-shifting prisms (right PA) biases attention to the left and is a promising technique for improving visuospatial neglect after right hemisphere damage (Rossetti et al., 1998; Azouvi et al., 2017). Right PA in neglect patients ameliorates not only visual abnormalities, such as the rightward shift in line bisection performance, but also extinction of tactile and auditory stimuli on the left, altered perception of time (for a review see Clarke and Crottaz-Herbette, 2016) and impaired mental time travelling (Anelli, n.d.). Left PA produces neglect-like behavior in healthy individuals (Colent et al., 2000; Schintu et al., 2014) and affects not only visuospatial cognition, but also spatial remapping, perception of time (for a review see Michel, 2016), and feedback learning performance (Schintu et al., 2018).

There is general agreement regarding the behavioral consequences of PA. However, the underlying neural changes are not clear and the results of the few studies investigating them are inconsistent. Functional imaging studies of PA have implicated the cerebellum (Weiner et al., 1983; Küper et al., 2013) and posterior parietal areas during both adaptation (Clower et al., 1996; Luauté et al., 2009; Chapman et al., 2010) and the aftereffect phase (Crottaz-Herbette et al., 2014; Magnani et al., 2014; Schintu et al., 2016). There is also evidence that PA affects behavior by acting on the dorsal frontoparietal network (Striemer and Danckert, 2010; Saj et al., 2013; Magnani et al., 2014; Schintu et al., 2016), which controls visually guided motor behavior and visual attention (Corbetta and Shulman, 2002; Milner and Goodale, 2006). However, while some studies have found only frontoparietal network involvement (Saj et al., 2013), others (Luaute et al., 2006; Crottaz-Herbette et al., 2017) describe changes extending to the ventral attentional network in the temporal lobe. Whether PA affects both the dorsal and ventral networks is still unknown. Another open question is whether the PA-induced changes are bilateral or unilateral, and, if bilateral, whether the direction of change in function is the same in both hemispheres. According to an influential model (Pisella et al., 2006; Striemer and Danckert, 2010) right PA induces its leftward attentional shift in neglect patients by decreasing activity in the intact left hemisphere and increasing it in the right, whereas left PA causes rightward bias in healthy individuals by doing the opposite. A few studies (Luaute et al., 2006; Crottaz-Herbette et al., 2014, 2017) support this model, but others favor a unilateral (Tsujimoto et al., 2018) or a bilateral and unidirectional (Saj et al., 2013) effect.

One reason for the inconsistencies in the literature could be that most functional imaging studies of PA have employed event-related designs and measured local changes in activity. Task-related activation changes may fail to identify brain areas whose connectivity is affected by PA but are not activated by the task. The task itself may also alter the state of the visual attention system in ways which obscure the effects of PA. Resting-state functional connectivity (RSFC) measures dynamic changes in connectivity within networks, without depending on task-related activation. In the one existing study of RSFC changes following PA (Tsujimoto et al., 2018), right PA modulated RSFC in the right dorsal network. However, the analysis was limited to regions of interest in the dorsal and ventral networks and may have failed to reveal changes in other areas. Investigating the effect of PA on RSFC across the entire brain, without choosing networks *a priori*, is a more rigorous test of mechanistic hypotheses and may reveal previously hidden aspects of the PA mechanism, helping to reconcile conflicting results in the literature.

The aim of this study was to investigate the effect of PA at the whole brain level by comparing the effects of left and right PA on RSFC in healthy subjects. Based on the prevailing PA model (Pisella et al., 2006; Striemer and Danckert, 2010) we hypothesized that PA should differentially change behavior and RSFC according to the direction of the visual displacement: Right PA should produce a leftward behavioral bias in association with decreasing RSFC in the left frontoparietal network and increasing RSFC in the right, whereas left PA should induce a rightward behavioral bias with opposite connectivity changes.

## MATERIAL AND METHODS

### Participants

Forty adults, free of neurological disorders or medications affecting brain function participated in the study. All had normal or corrected-to-normal vision, were right-handed (Edinburgh Inventory; Oldfield, 1971), and were right-eye dominant (hole-in-card test; Miles, 1930). Participants were compensated for participation and gave written informed consent. The study was approved by the National Institutes of Health, Central Nervous System Institutional Review Board, and conducted in accordance with the ethical standards of the 1964 Declaration of Helsinki (World Medical Association, 2013).

Twenty participants (12 female; age = 26.25 ± 0.87 SEM) underwent left PA and the other twenty, (13 female; age = 26.12 ± 1.05 SEM) right PA. After data collection, two participants were excluded from the right PA group, one because excessive motion during scans (average motion > 0.2 cm) and one because of a congenital cerebral cyst, which might have been associated with cortical reorganization. The data submitted to the statistical analysis were gathered from a total of 38 participants: left PA group (12 female; age = 26.25 ± 0.87 SEM) and right PA group (12 female; age = 25.73 ± 1.09 SEM). The left and right PA groups did not differ in age (t(36) = 0.375, p = .710).

## Procedure

The experiment consisted of two sessions of behavioral testing and fMRI, one before, and one after, PA (Figure 1). In each session, we measured visual perception with the perceptual line bisection and manual line bisection tasks. Then participants underwent neuroimaging consisting of resting state scan and population receptive field scans (pRF; Wandell et al., 2007, which is part of a future report and not discussed further here). Following the resting state and pRF scans, we repeated the perceptual line bisection and manual line bisection tasks, along with two tasks assessing proprioceptive (straight-ahead pointing) and sensorimotor (open-loop pointing) performance (Figure1). Participants then received left or right PA. Immediately after PA (early post adaptation assessment), we assessed proprioceptive and sensorimotor performance with the straight-ahead and open-loop pointing tasks, and performed another resting state and pRF scan, followed by the perceptual and manual line bisection and the straight-ahead and open-loop pointing tasks (late post adaptation assessment).

**Figure 1.**
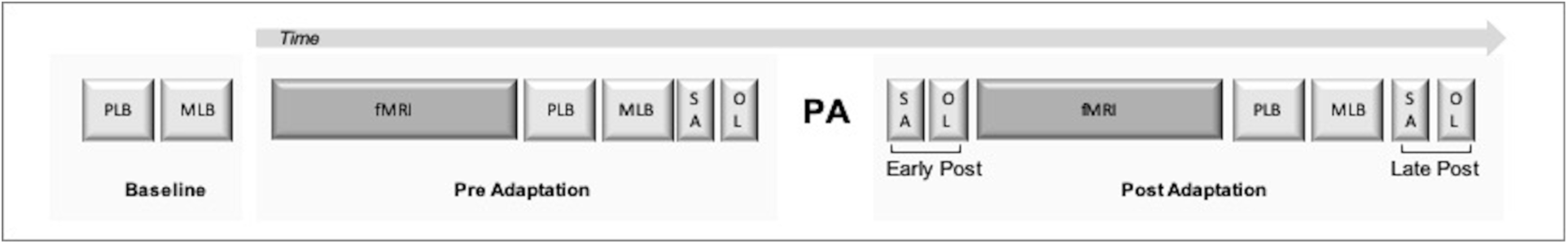
Experimental Design. PLB = perceptual line bisection; MLB = manual line bisection; OL = open-loop pointing; SA = straight-ahead pointing; fMRI = functional magnetic resonance imaging, PA = prism adaptation.

During the behavioral measures and prism adaptation, participants were seated in front of a horizontal board with their heads supported by a chinrest. On the board, three circular targets (8 mm in diameter) were positioned at 0, −10 (left), and +10 (right), degrees from the body midline, approximately 57 cm from participant’s nasion and were used for the prism adaptation, open-loop and straight-ahead pointing tasks.

### Prism Adaptation

During PA, participants were fitted with prism goggles with a 15° left (left PA) or right (right PA) visual field deviation and performed 150 pointing movements to the right and left targets in a verbally cued, pseudorandom, order. Before each pointing movement, participants placed their right index finger in the starting position on a 1.5 cm diameter pad, located close to the midline of the chest. Participants could not see their hands in the starting position and during the first third of the pointing movement. Participants were instructed to point with the index finger extended, in a single movement at a fast but comfortable speed, and to return the hand to the starting position.

### Behavioral assessment

#### Perceptual line bisection

prioritizes the perceptual, and minimizes the motor, component of the visuospatial bias by asking participants to judge a series or pre-bisected lines instead of actively bisecting them. We used a modified version of the Landmark task (Milner et al., 1992) The task consisted of 66 white, pre-bisected, lines (350 mm x ∼2 mm) displayed on a black screen positioned 35 cm from the eyes. Lines were transected at the true center and at 2, 4, 6, 8, and 10 mm to the left and right of the true center. Each of the 11 different pre-bisected lines was presented six times in a pseudorandom order, yielding a total of 66 trials, which took approximately three minutes to complete. Each line was displayed for a maximum of five seconds or until a response was made, and was then replaced by a black-and-white, patterned mask, which stayed on the screen for one second before the next line was displayed. We used Presentation software (Neurobehavioral Systems, Inc., USA) to generate the stimuli, record responses, and control the task. Participants were instructed to inspect each line and judge whether the transecting mark was closer to the left or right end and to respond within 5 seconds by pressing pedals positioned under the left and right feet. We chose a pedal response to limit the use of the right hand, which was used for PA, since post-adaptation feedback from that hand could contribute to de-adaptation. Subjects performed at least ten practice trials before the baseline measurement. For each participant, we plotted the percentage of right-side responses as a function of the position of the transector (true center and 2,4,6,8,10 mm to left and to the right of the true center). We then fit a sigmoid function to the data. The value on the x-axis corresponding to the point at which the participant responded with the right pedal 50% of the time was taken as the point of subjective equality (PSE).

#### Manual line bisection

emphasizes the motor over the perceptual component of the visuospatial bias (Milner et al., 1992). We used this task to measure the visuospatial shift induced by PA (Schenkenberg et al., 1980). It consisted of a series of 10 black lines (identical in size to those used for the perceptual line bisection task) drawn on 297mm × 420mm sheets of paper, which were positioned over the same computer screen. Participants were instructed to inspect each line and, with a pen held in their right hand, draw a vertical mark at the perceived center of each line. No time limit was imposed, and participants took on average of 1 second to place the mark on each line. We measured the distance between the mark placed by the participant and the true center of the line and took the average as the PSE, with marks to the right of center coded as positive.

#### Straight-ahead pointing

was used to measure the proprioceptive shift induced by PA. Participants performed six pointing movements to the midline with the right index finger at a comfortable and uniform speed, while resting their left hands on their laps. Before each movement, participants were told to close their eyes and imagine a line splitting their body in half and to project it onto the board in front of them. We then asked them to point to the line with their eyes closed and return to the starting position. To ensure that participants had no visual feedback, the arm and hand were occluded by a cardboard baffle before movement onset. The proprioceptive shift was measured as the average distance between the landing position and the true midline with precision of +/-0.5 cm.

#### Open-loop pointing

was used to measure the sensorimotor shift induced by PA. Participants performed six pointing movements with the right index finger to the central (0°) target at a comfortable and uniform speed, while resting their left hands on their laps. The experimenter noted the landing position of the participant’s finger with a precision of +/-0.5 cm. Before each movement, we instructed participants to look at the central target, close their eyes, point to the target while keeping their eyes closed, and then return the hand to the starting position. As in the straight-ahead task, vision of the arm and hand was occluded. We measured the sensorimotor shift as the average distance between the landing position and the central target.

### fMRI

#### MRI procedure

We acquired functional and structural MRI data with a 32-channel head coil on a research-dedicated 3-Tesla Siemens MAGNETOM Prisma MR scanner. Head movement was minimized with padding. A whole-brain T1-weighted anatomical image (MPRAGE) was obtained for each participant (208 slices, voxel size 1.0 × 1.0 × 1.0 mm, repetition time (TR) = 2530 ms, echo time (TE) = 3.3 ms, TI = 1100 ms, field of view (FOV) = 256 × 256 × 208 mm, flip angle = 7°). T2* blood oxygen level-dependent (BOLD) resting state scans were acquired for all subjects (46 slices aligned to the AC-PC axis, voxel size 3.0 × 3.0 × 3.0 mm, TR = 2500 ms, TE = 30.0 ms, FOV = 192 x 138 x 192 mm, flip angle 70°, 64 × 46 × 64 acquisition matrix). During resting state scans, lighting was dimmed, and subjects were instructed to lie still and look at a white central cross appearing on a black screen.

#### MRI preprocessing

Functional and structural MRI data were preprocessed using the AFNI (version 18.2.15) software package (Cox, 1996) and followed the general preprocessing approach of (Wang et al., 2014). The anatomical scans were segmented into tissue compartments using Freesurfer. We removed the two initial volumes from each resting state scans to allow the magnetic field to stabilize. Volumes were then despiked (3dDespike), slice-time corrected to the first slice, co-registered to the anatomical scan, transformed to TT_N27 Template space (Talairach and Tournoux 1988), resampled to 2 mm isotropic voxels, smoothed with an isometric 4-mm full-width half-maximum Gaussian kernel, and scaled to percentage signal change (dividing each voxel’s timeseries by its mean). TRs with head movement greater than 0.3 mm were censored from the analysis, simultaneously with bandpass filtering (using 3dTproject) from 0.01 to 0.1 Hz. We regressed the 6 motion parameters and their derivatives, which were also filtered in the same manner (0.01 to 0.1 Hz) prior to performing the nuisance regression (e.g., Hallquist et al., 2013; Jo et al., 2013). Measures of mean framewise displacement (using the AFNI function @1dDiffMag) and average voxelwise signal amplitude (standard deviation) were also calculated for use as nuisance covariates in group-level analyses in order to control for any residual global artifacts in the resting-state scans (Wang et al., 2014).

#### Functional connectivity analysis

To initiate the analysis, we created seed regions composed of the right and left intra-parietal sulcus (IPS) regions 1 and 2, where transcranial magnetic stimulation causes changes in visuospatial behavior when applied online (Szczepanski and Kastner, 2013). We identified these regions of interest using a probabilistic atlas of the visual areas (Wang et al., 2015). We created each seed by transforming the IPS 1 and 2 maps from the probabilistic atlas into TT_N27 template space and keeping voxels that had ≥ 30% probability of classification as being in IPS 1 or 2. We then combined the IPS 1 and 2 voxels to form a seed region (IPS1-2; Figure 3a). We created one seed in each hemisphere (left IPS1-2 volume = 2984mm^3^, right IPS1-2 volume = 2672mm^3^; Figure 3A) and whole-brain time series correlation maps from each seed (Pearson correlation followed by Fisher’s z-transform to improve normality). After functional MRI preprocessing, we created a brain mask for each participant, which included voxels with functional data present and excluding non-neural tissue (e.g., ventricles, white matter). We then created a group-level brain mask from these individual masks for use in group analyses using voxels where at least 90% of participants had data.

**Figure 2.**
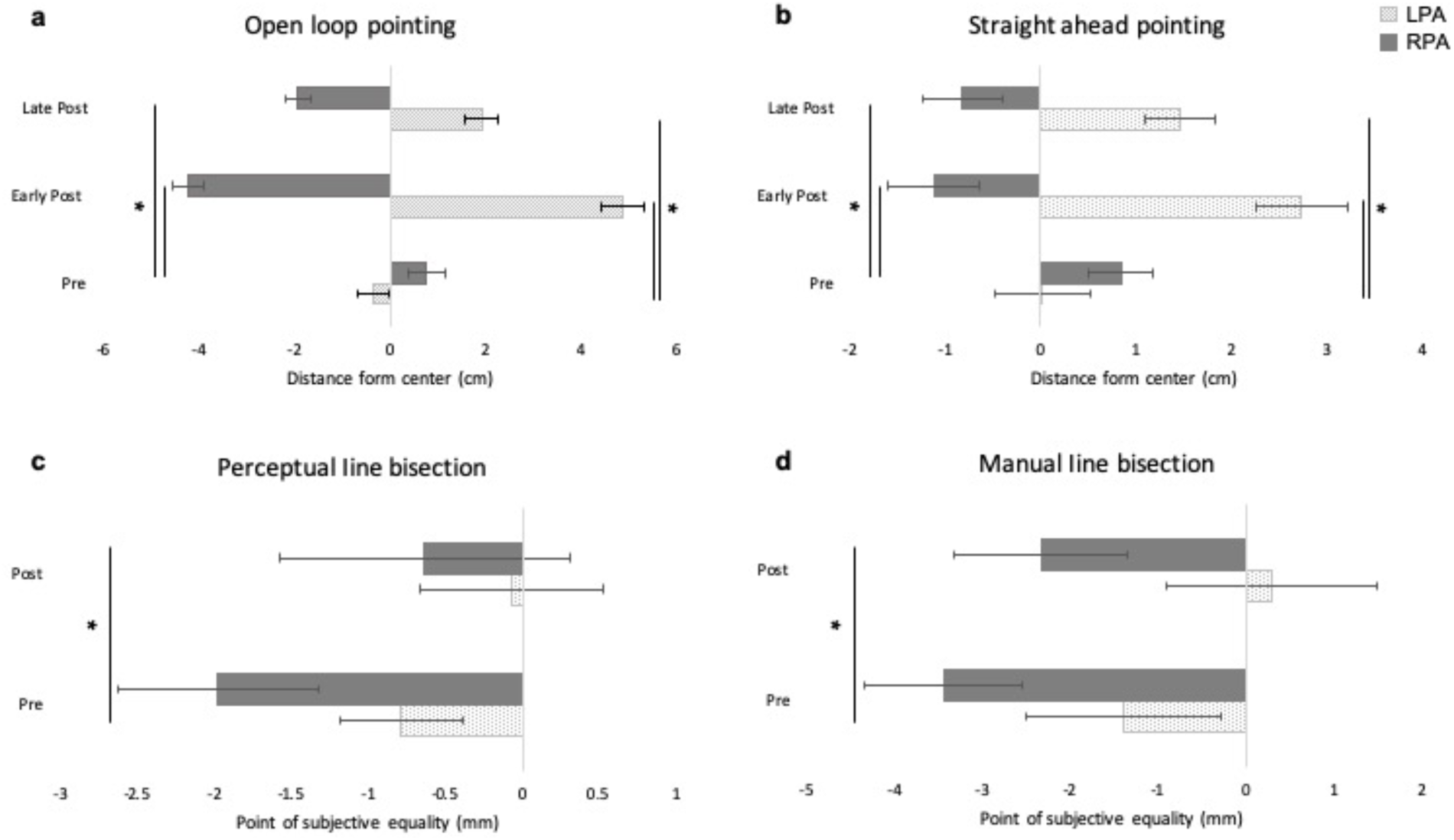
Behavioral effects of prism adaptation. Negative and positive values represent left and right of center, respectively. Error bars represent 1 SEM.

**Figure 3.**
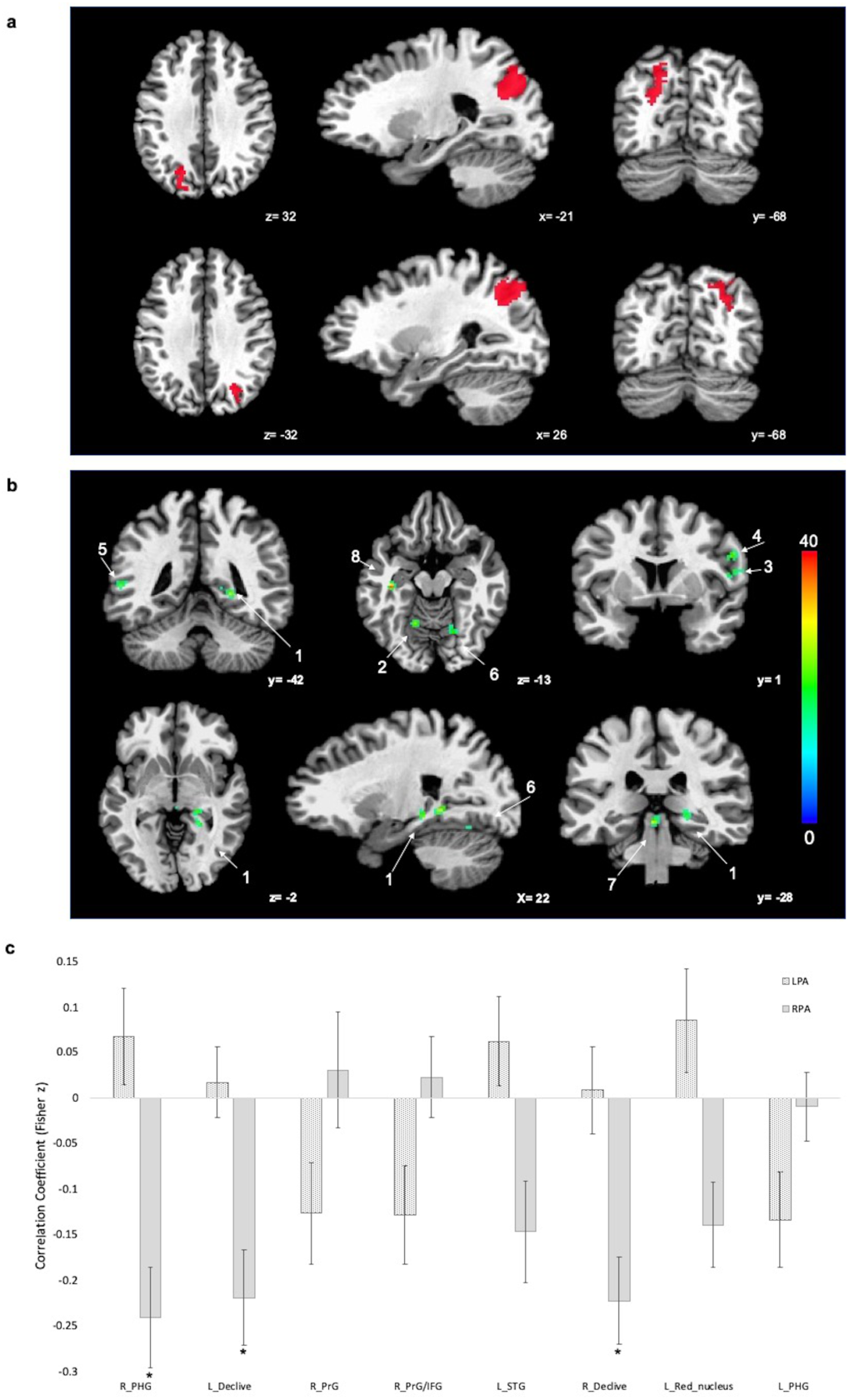
**a:** Locations of left and right IPS1-2 seeds **b**: Brain regions with significant clusters of RSFC change resulting from the Group x Time interaction (false discovery rate (FDR) corrected, p < 0.05). Color scale indicates F-value. **c:** Amount of change (post *minus* pre) in RSFC between each cluster in A and the IPS1-2 seeds. Error bars represent 1 SEM. *p < .001 and surviving FDR-correction. Only clusters of 20 or more adjacent voxels were retained.

### Statistical analysis

Statistical analyses were performed using SPSS (IBM, Version 24.0), R and Matlab (R2016a), AFNI (3dLME command) with family-wise alpha set at .05. All data are presented as mean and the standard error (SEM). Effect sizes were computed as Cohen’s d. When sphericity was violated, Greenhouse-Geissser corrected values are reported. We used paired or independent t-tests for post-hoc comparisons. To assess the changes in RSFC, data were submitted to a linear mixed effects regression model (LMER) with seed-based functional connectivity (correlation maps) as the dependent variable, Group (left PA, right PA), Time (pre, post), and Hemisphere (left, right) as fixed effects, Subject as a random effect, and motion (@1dDiffMag) and average voxel-wise standard deviation as nuisance covariates. Pearson correlation was computed in AFNI (3dTcorr1D command) between the neural (whole-brain correlation maps, post *minus* pre) and behavioral amount of change (post *minus* pre) with FDR correction to *q*=0.05.

## RESULTS

### Behavior

#### Open-loop pointing

We measured sensorimotor performance by quantifying the deviation in pointing from the landing position and the true center, before (pre) immediately after prism adaptation (early-post), and at the end of the experiment (late-post). The mixed analysis of variance (ANOVA) with Time (pre, early-post, late-post) as a within-participant variable and Group (left PA, right PA) as a between-participant variable revealed a significant main effect of Group (F(1, 36) = 78.363, p = 0.002, η^*2*^_*p*_*=* 0.68), such that the pointing error was right of the center (mean = 2.164 cm) for the left PA group and left of the center (−1.795 cm) for the right PA group (Figure 2a). There was a significant Time x Group interaction (F(2, 36) = 284.031, p < 0.001, η^*2*^_*p*_*=* 0.89) and non-significant main effect of Time (F(2, 36) = 1.169, p = 0.317). Follow-up repeated-measures ANOVAs, with Time as a within-participants variable, performed individually for each group to unpack the significant Time x Group interaction, revealed a main effect of Time for both left PA (F(2, 19) 193.373 p < 0.001, *η*^2^_p_= 0.91) and right PA (F(2,17) = 106.951, p < 0.001, *η*^2^_p_= 0.86) groups. The left PA group showed a rightward shift in pointing at both the early (4.86 cm) (t(19) = −17.228, p < 0.001, Cohen’s d = −3.85) and late (1.92 cm) (t(19) = −9.703, p < 0.001, Cohen’s d = −2.17) post measurements. The right PA group exhibited a leftward shift at the early post (−4.22 cm) (t(17) = −12.196, p < 0.001, Cohen’s d = 2.73) and late post (−1.93 cm) (t(17) = 9.694, p < 0.001, Cohen’s d = 2.17) measurements. Independent t-tests on the absolute values of change found no significant difference between the groups in the size of the shift at either the early (t(36) = 0.470, p = 0.641) or late (t(36) = - 1.137, p = 0.263) post measurements.

#### Straight-ahead pointing

We measured proprioceptive performance by quantifying the deviation in the pointing between the perceived and true participants’ midline. The mixed effects ANOVA with Time (pre, early-post, late-post) as a within-participants variable and Group (left PA, right PA) as a between-participants variable revealed a significant main effect of Group (F(1, 36) = 11.661, p = 0.002, η^*2*^_*p*_*=* 0.24; Figure 2b). Again, the pointing error was rightward for the left PA group (1.404 cm) and leftward for the right PA group (−0.363 cm). There was a significant Time x Group interaction (F(2,36) = 33.754, p < 0.001 η^*2*^_*p*_*=* 0.48) and a non-significant main effect of Time (F(2, 36) = 1.534, p = 0.223). Follow-up repeated measures ANOVAs performed individually for each group found a significant main effect of Time for both left PA (F(2,19) = 27.669, p < 0.001, η^*2*^_*p*_*=* 0.59) and right PA (F(2,17) = 10.729, p < 0.001, η^*2*^_*p*_*=* 0.39). The left PA group had a rightward shift in pointing at both the early (2.733 cm) (t(19) = −6.176, p < 0.001, Cohen’s d = −1.38) and late (−1.458) (t(19) = −4.604, p < 0.001, Cohen’s d = −1.03) post measurements, and right PA group pointing shifted leftward; at the early (−1.120 cm) (t(17) = 4.182, p = 0.001, Cohen’s d = 0.93) and late (−0.815 cm) (t(1, 17) = 4.880, p < 0.001, Cohen’s d = 1.09) time point. Independent t-tests comparing the absolute value of change revealed no significant difference in the aftereffect between the two groups at early (t(36) = 0.676, p = 0.503) and late (t(36) = −0.510, p = 0.613) post measurements.

#### Perceptual line bisection

We averaged the two pre-adaptation scores, since they did not differ (t(36) = 1.585, p = 0.122). The mixed ANOVA with Time (pre, post) as within-participants variable and Group (left PA, right PA) as between-participants variable revealed a significant main effect of Time (F(1, 36) = 5.363, p = 0.026, η^*2*^_*p*_*=*0.13; Figure 2c). Both groups shifted their midline judgment rightward (from −1.317 mm to - 0.338 mm) after PA. The Time x Group interaction was not significant (F(1, 36) = 0.491, p = 0.488).

#### Manual line bisection

As for the perceptual line bisection task, we averaged the two pre-adaptation scores since they did not differ (t(36) = 1.418 p = 0.165). A mixed effects ANOVA with Time (pre, post) as a within-participants variable and Group (left PA, right PA) as a between-participants variable, showed a main effect of Time (F(1, 36) = 9.101, p = 0.005, η^*2*^_*p*_*=* 0.202). PA produced a rightward bias in midline judgment independent of PA direction (from −2.365 mm to −0.958 mm). The Time x Group interaction was not significant (F(1, 36) = 0.378, p = 0.543).

### Resting State Functional Connectivity

Before covarying head motion and the average standard deviation nuisance measures in the LMER, we ascertained that those variables did not differ between the RPA and LPA groups at baseline (motion t(36) = -.119, p = .906; standard deviation t(36) = 1.004, p = .322; RPA: motion mean = .062 (SEM = .008), standard deviation = .826 (0.110); LPA: motion = .064 (.007), standard deviation = .785 (0.141)) or at post (motion t(36) = .129, p = .898; standard deviation t(36) = -.434, p = .667; RPA: motion = .060 (.008), standard deviation = .790 (.021); LPA: motion .059 (.005), standard deviation = .806 (.028)). Similarly, the within comparison did not reveal any difference between pre- and post-PA phases for both the RPA (motion t(17) = -.276, p = .786; standard deviation t(17) = −1.503, p = .151) and LPA groups (motion t(19) = −1.041, p = .311; standard deviation t(19) = .962, p = .348).

The LMER analysis revealed a significant Group x Time interaction (FDR-corrected to p < 0.05; q < 0.05), such that RSFC was differentially affected according to PA direction. The analysis detected 8 significant clusters relative to the IPS seeds (Table 1 and Figure 3b). We included any clusters with 20 or more voxels in subsequent analyses in order to avoid post-hoc testing in small, noisy clusters.

**Table 1.** Clusters surviving the Group x Time interaction (FDR corrected q < 0.05; p < .05). Coordinates are in Talairach-Tournoux space.

| Volume<br>(mm<sup>3</sup>) | Peak<br>(X) | Peak<br>(y) | Peak<br>(z) | Brain Region |
| --- | --- | --- | --- | --- |
| 520 | 21 | -43 | 0 | Right Parahippocampal Gyrus |
| 288 | -17 | -55 | -14 | Left Declive |
| 264 | 53 | -3 | 14 | Right Precentral Gyrus |
| 200 | 53 | 3 | 28 | Right Precentral Gyrus/Inferior Frontal Gyrus |
| 192 | -55 | -43 | 6 | Left Superior Temporal Gyrus |
| 184 | 17 | -63 | -12 | Right Declive |
| 176 | -3 | -29 | -8 | (Within 6 mm) Left Red nucleus |
| 168 | -37 | -23 | -14 | Left Parahippocampal Gyrus |

To follow up the Group x Time interaction, we extracted the time series of each cluster that survived the LMER for each group separately. We computed correlations between each cluster time series and the averaged time series of the IPS1-2 seeds. We then compared the Fisher z-transformed correlation coefficient before and after PA. Post-hoc testing (paired t-tests) of the correlation coefficients revealed that the Time x Group interaction was characterized mainly by a decrease in RSFC between the IPS seeds and other brain regions, and that the changes were specific to the PA direction, as shown in Figure 3c.

Right and left PA had differential effects on RSFC between the IPS seeds and the parahippocampal gyri. Right PA caused a decrease in RSFC between the seeds and a cluster including the posterior portion of the right parahippocampal gyrus, the fusiform and lingual gyri, and extending to the thalamus (t(17) = 4.367, p < 0.001, Cohen’s d = 1.03). The left PA group showed a decrease in connectivity between the IPS seeds and a cluster including the anterior portion of the left parahippocampal gyrus and extending to the hippocampus (t(19) = 2.531, p = 0.02, Cohen’s d = 0.57). The right PA group also showed changes in FC between the IPS seeds and the superior temporal gyrus (t(17) = 2.617, p = 0.018, Cohen’s d = 0.62), red nucleus (t(17) = 2.939, p = 0.009, Cohen’s d = 0.569), the left (t(17) = 4.176, p = 0.001, Cohen’s d = 0.98) and right (t(17) = 4.646, p < 0.001, Cohen’s d = 1.09) cerebellum. By contrast, the left PA group showed a decrease in FC between the seeds and the right precentral (t(19) = 2.252, p = 0.036, Cohen’s d = 0.50) and inferior frontal gyri (t(19) = 2.360, p = 0.029, Cohen’s d = 0.53). Following FDR correction (q < .05, p-value threshold = 0.009), only the following clusters survived and only in the right PA group: fusiform and lingual gyri and extending to the thalamus (t(17) = 4.367, p < 0.001, Cohen’s d = 1.03), and the left (t(17) = 4.176, p = 0.001, Cohen’s d = 0.98) and right (t(17) = 4.646, p < 0.001, Cohen’s d = 1.09) cerebellum.

There were differential changes in RSFC between the IPS seeds and other brain structures, which did not survive the whole brain analysis, but did survive the single group contrasts, corrected for whole-brain contrasts (FDR corrected q < .05; voxelwise threshold p = .0051). As reported in Table 2, the separate contrasts (post versus pre) for each group revealed that both left and right PA decreased RSFC between IPS seeds and the superior temporal gyrus (STG) bilaterally (Figure 4a1, Figure 4a2, Figure 4b1, Figure 4b2). Right PA increased RSFC between the IPS seeds and the right middle frontal gyrus (MFG; Figure 4a3), and left PA decreased it (Figure 4b3). Right PA increased RSFC between the IPS seeds and inferior parietal lobule bilaterally (IPL; Figure 4a4, Figure 4a5), and left PA decreased RSFC between the IPS seeds and the left IPL (Figure 4b4), and between the IPS seeds and the right superior parietal lobule (SPL; Figure 4b5). See Tables 4 (right PA) and 5 (left PA) in the supplementary material for a full list of significant clusters).

**Table 2.**
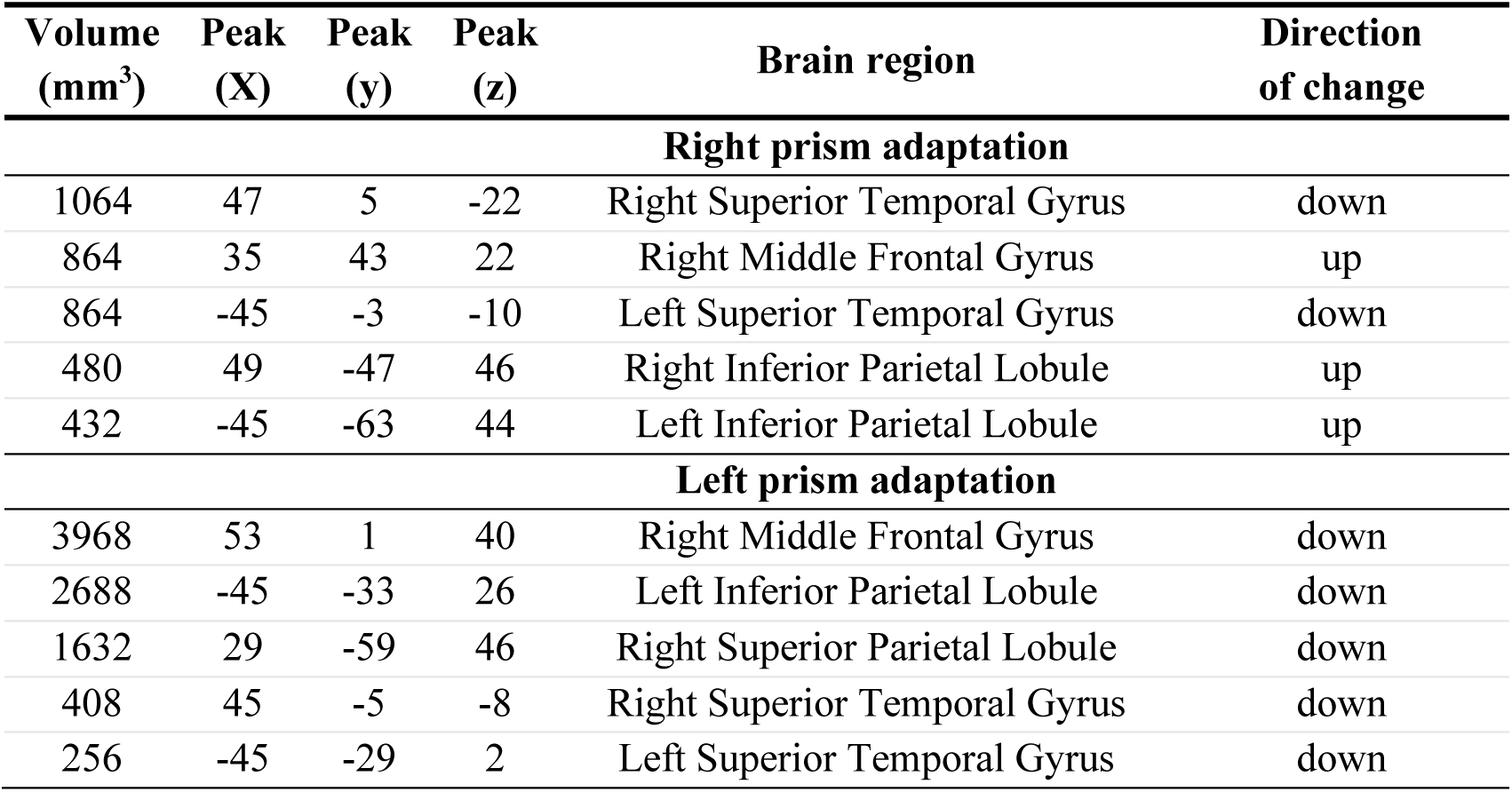
Contrast (post - pre) for right PA and left PA. Coordinates are in Talairach-Tournoux space. BA = Brodmann Area.

| Volume<br>(mm<sup>3</sup>) | Peak<br>(X) | Peak<br>(y) | Peak<br>(z) | Brain region | Direction<br>of change |
| --- | --- | --- | --- | --- | --- |
| **Right prism adaptation** |  |  |  |  |  |
| 1064 | 47 | 5 | -22 | Right Superior Temporal Gyrus | down |
| 864 | 35 | 43 | 22 | Right Middle Frontal Gyrus | up |
| 864 | -45 | -3 | -10 | Left Superior Temporal Gyrus | down |
| 480 | 49 | -47 | 46 | Right Inferior Parietal Lobule | up |
| 432 | -45 | -63 | 44 | Left Inferior Parietal Lobule | up |
| **Left prism adaptation** |  |  |  |  |  |
| 3968 | 53 | 1 | 40 | Right Middle Frontal Gyrus | down |
| 2688 | -45 | -33 | 26 | Left Inferior Parietal Lobule | down |
| 1632 | 29 | -59 | 46 | Right Superior Parietal Lobule | down |
| 408 | 45 | -5 | -8 | Right Superior Temporal Gyrus | down |
| 256 | -45 | -29 | 2 | Left Superior Temporal Gyrus | down |

**Figure 4.**
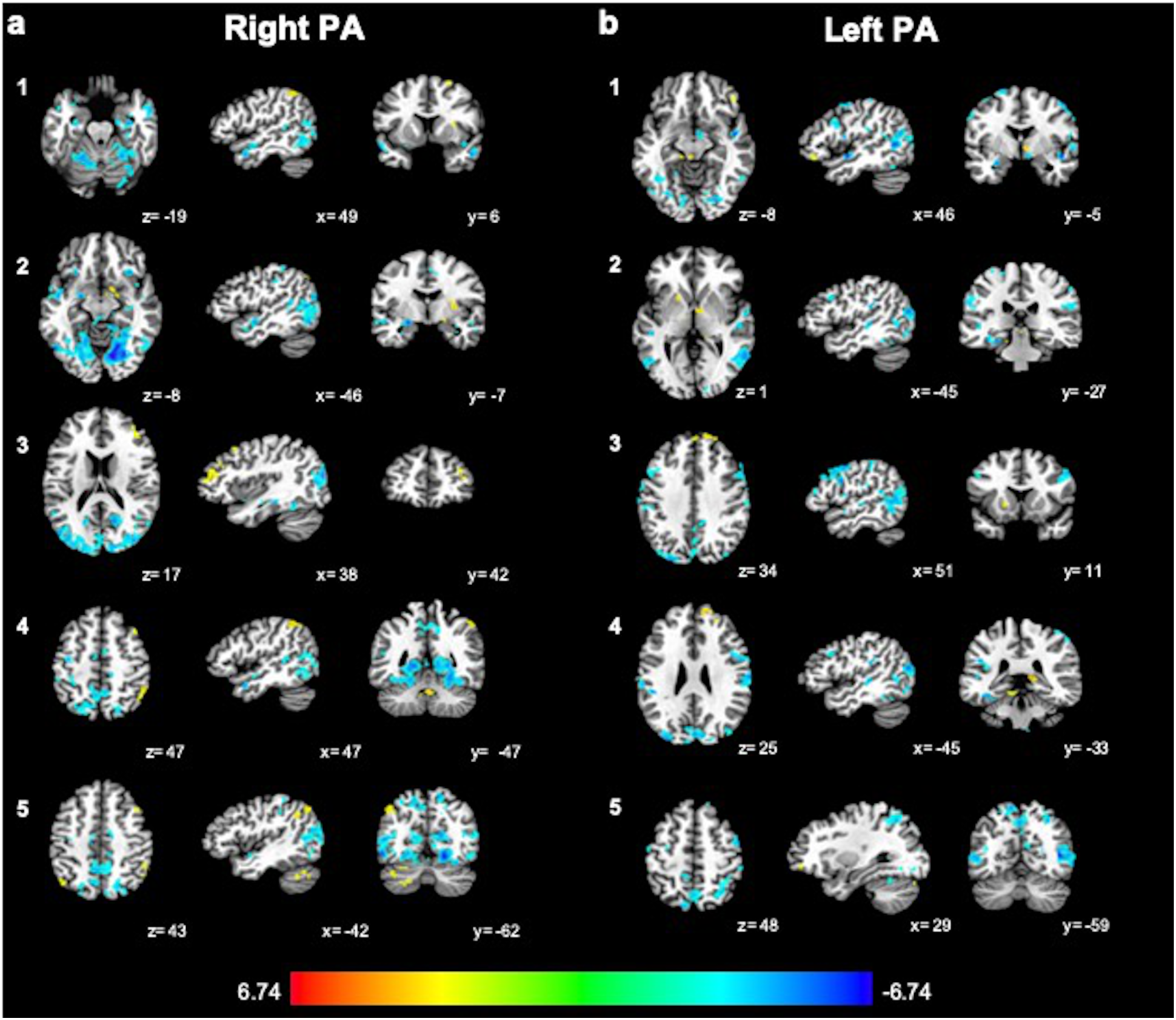
Contrast (post vs. pre) z-score for the right (a) and left (b) PA groups, FDR-corrected for multiple comparisons (FDR q<.05, p < 0.0052). Color scale indicates z-value, with warm colors indicating RSFC increases with IPS1-2 and cool colors indicating RSFC decreases.

Finally, to examine whether neural changes in the right hemisphere following PA were directly related to the corresponding behavioral changes for the RPA group we computed a Pearson correlation between the amount of change (post *minus* pre) in RSFC between right IPS 1-2 and the rest of the brain (whole-brain correlation maps) and the amount of change (post *minus* pre) of the performance at the two pointing task as those where the two behavioral tasks that showed significant opposite effects according to PA direction. The analysis revealed a significant correlation surviving whole-brain comparisons between the behavioral changes at the straight-ahead pointing task (p<.0005, q = .05) and not in the open-loop pointing task (see Supplementary Table 3 for a full report).

## DISCUSSION

We investigated the effect PA on RSFC at the whole brain level by contrasting the effects of left and right PA in two groups of healthy participants and found differential changes in a cerebellar-parieto-parahippocampal network in the right hemisphere, depending on the direction of prism adaptation. We also found that left and right PA induced opposite changes in RSFC between regions within both PPCs and between the frontal and parietal areas involved in visuospatial function. We also confirmed the involvement of the superior temporal area in the neural response to PA.

### Behavioral results

As expected, left PA produced a rightward shift, and right PA produced a leftward shift, in sensorimotor and proprioceptive pointing performance. These aftereffects were of similar magnitude, so any differences in the imaging results cannot be attributed to a difference in the magnitude of the adaptation. The aftereffects were still present at the end of the experiment, meaning that the subjects were still adapted while fMRI data were acquired and visuospatial behavior assessed.

While we expected a significant rightward bias (neglect-like behavior) on the perceptual line bisection task after left, but not right, PA (Colent et al., 2000), there was a significant rightward shift in midline judgment, independent of PA direction, on both the perceptual and manual line bisection tasks. However, this was not surprising, since the behavioral effects of right PA in healthy individuals are inconsistent (Schintu et al., 2017) and previous work has reported a rightward aftereffect with right PA (Berberovic and Mattingley, 2003).

### Neuroimaging results

Compared to left PA, right PA decreased RSFC between a portion of the PPC (the IPS1-2 seeds) and the parahippocampal gyrus, both of which are engaged during spatial navigation (Aguirre et al., 1996; Aguirre and D’Esposito, 1997; Maguire et al., 1998; Grön et al., 2000). The PPC, which contains egocentric (body-referenced) representations (Silver and Kastner, 2009), may feed spatial information to the parahippocampal cortex, which is important for allocentric (world-referenced) representation (Aguirre and D’Esposito, 1999). The PPC appears to transform the allocentric output of the hippocampus and other medial temporal lobe structures into egocentric coordinates to support movement through the environment (Whitlock et al., 2008; Kravitz et al., 2011). RSFC between a portion of the PPC and parahippocampal gyrus is decreased following right PA, suggesting that reduced communication between these areas could weaken the influence of the egocentric reference frame in the navigation network, making it more dependent on allocentric information. The reduction of connectivity between PPC and cerebellum after right PA could further decrease the contribution of the egocentric reference frame to navigation-related regions. The cerebellum is not only involved in spatial navigation (Malm et al., 1998; Schmahmann and Sherman, 1998; Molinari et al., 2004), but linked with brainstem and thalamic structures concerned with oculomotor control and the vestibular system (Schmahmann, 2010), which encodes self-orientation with respect to gravity.

The cerebellum may transform the reference frame of vestibular signals and convey them to the hippocampus via a multi-synaptic pathway involving the PPC (Rochefort et al., 2013). As the model of PA proposes (Pisella et al., 2006), and given the connection with the PPC (Casula et al., 2016), it is possible that the cerebellum supplies egocentric spatial data to the PPC via bottom-up signals. The cerebellum is known to be involved in visuo-motor transformation learning (Graydon et al., 2005), which is required for the PA adaptation phase. However, we found that the changes in RSFC between cerebellum and PPC outlasted adaptation and persisted in the aftereffect phase.

Right and left PA have different effects on RSFC between the PPC and parahippocampal cortex and between the PPC and cerebellum, characterized as the “navigation network” (Maguire et al., 1998). Therefore, right PA, by downregulating connectivity between the PPC and the cerebellum, may reduce vestibular input, and, by decreasing it between the PPC and right parahippocampal gyrus, further decrease the influence of egocentric spatial information in favor of allocentric signaling. This effect on the spatial reference frame may explain how right PA improves the complex of symptoms in neglect. Vestibular input is important for maintaining spatial orientation (Ventre-Dominey et al., 1999; Doricchi et al., 2002) and plays a role in neglect (Cappa et al., 1987; Vallar et al., 1993). For example, irrigation of the right ear with cold water, to cause convection in the endolymph of the vestibular apparatus and stimulate vestibular signaling, improves spatial functioning in neglect patients. Vestibular stimulation, especially when combined with activation of neck muscle stretch receptors, another source of positional information, by vibration, provides the sensory signals needed to create a spatial frame of reference based on eye and head position in space (Karnath, 1994). Eye and head position perception are compromised in neglect. Since neglect patients have impaired vestibular signaling to the IPS, egocentric spatial information relayed to the PPC may be inaccurate. Right PA could decrease those erroneous inputs in favor of unimpaired allocentric ones, partially restoring spatial perception. This interpretation is supported by the correlation between the amount of change in RSFC between right IPS 1-2 and the rest of the brain, and the degree of behavioral adaptation measured by the straight-ahead pointing task, which is mainly based on the egocentric reference frame.

The contrast between left and right PA revealed those brain regions whose RSFC changes differed between the two PA directions and survived whole brain correction. However, post hoc analysis allowed us to look for changes in RSFC before and after PA within each group separately. At the single group level, right PA increased RSFC between areas within the PPCs, whereas left PA decreased it. The bilateral effect on local connectivity within the PPC is consistent with previous findings (Saj et al., 2013) showing a bilateral increase of task-related activation in parietal areas. Right PA increased, and left PA decreased, RSFC between the seeded portion of the PPC and the MFG. The posterior IPS is the node of the dorsal attentional network receiving the most input from the ventral attentional network, possibly via the right MFG, which is thought to be the link between the two networks (Corbetta et al., 2008). Activity in the right MFG correlates with activity of both attention networks (Fox et al., 2006) and disconnection in the right ventral attention network is strongly related to the severity of neglect (He et al., 2007). This is consistent with the effect of PA on frontoparietal RSFC in our data. Finally, both left and right PA decreased RSFC between the seed in the PPC and the STG bilaterally. The STG, where damage causes neglect (Karnath et al., 2001, 2004), is involved in visual attention, and has been affected by PA in previous studies (Luaute et al., 2006; Crottaz-Herbette et al., 2017).

In conclusion, we propose that right, compared to left, PA decreases egocentrically referenced input to the right parieto-cerebellar-parahippocampal navigation network (Maguire et al., 1998). Our results confirm the action of PA on the frontoparietal network, but only partially agree with the preexisting model of PA (Pisella et al., 2006; Striemer and Danckert, 2010). Our data confirm that PA, depending on direction of visual displacement, changes RSFC between areas within the PPCs and between the PPCs and the right frontoparietal network. However, they also refine the model by adding the knowledge that the direction of RSFC change is the same for both hemispheres and includes the STG whose connectivity with the PPCs decreases. Right PA may ameliorate visuospatial impairments in neglect by increasing RSFC between the IPS and frontoparietal cortex in the right, lesioned, hemisphere, and left PA may induce neglect-like behavior in the healthy by decreasing frontoparietal RSFC on the right, in both cases decreasing the influence of temporal over posterior parietal regions.

## Acknowledgment

This work was supported by the National Science Foundation (grant number BCS-1534823 to Prof. Shomstein), the National Institutes of Health Ruth L. Kirschstein National Research Service Award (to Dr. Schintu), the Center for Neuroscience and Regenerative Medicine (grant number CNRM-70-3904 to Dr. Freedberg), the Intramural Research Program of the National Institute of Mental Health (to Dr. Gotts), and the Clinical Neurosciences Program of the National Institute of Neurological Disorders and Stroke.

Selene Schintu, Ph.D. Postdoctoral fellow, Behavioral Neurology Unit, NINDS, NIH. 10 Center Drive, MSC 1440, Bethesda, MD 20892-1430

