## Supplemental for "Prism adaptation modulates connectivity of the intraparietal sulcus with multiple brain networks"

**Supplementary material**

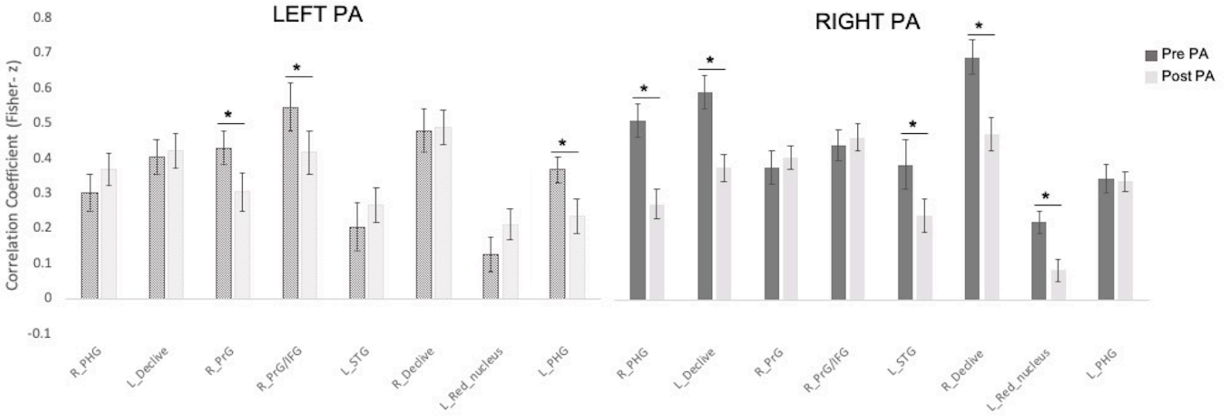


**Figure 5.** Change in connectivity (post *minus* pre) between each cluster and the average seeds (left and right IPS1-2 seeds). Error bars represent 1 SEM.

**Table 3**. Clusters surviving the Pearson correlation between the amount of change (post *minus* pre) in performance at the straight-ahead pointing task and changes in RSFC between right IPS 1-2 and the rest of the brain ($p < .0005$; $q = 0.05$). Only clusters of 20 or more adjacent voxels were retained. Coordinates are in Talairach-Tournoux space.

| **Volume ($mm^3$)** | **Peak (X)** | **Peak (y)** | **Peak (z)** | **Brain Region** |
|---|---|---|---|---|
| 352 | -29 | -45 | -24 | Left Culmen |
| 312 | 11 | -71 | -18 | Right Declive |
| 264 | -9 | -75 | 10 | Left Cuneus |
| 240 | -13 | -21 | 8 | Left Thalamus |
| 200 | -25 | -63 | -20 | Left Declive |
| 192 | -3 | 13 | 44 | Left Medial Frontal Gyrus |

**Table 4.** Contrast (post-pre) for the right PA. Coordinates are in Talairach-Tournoux space. BA = Brodmann Area. Only clusters of 20 or more adjacent voxels were retained.

| | Volume ($mm^3$) | Peak (X) | Peak (y) | Peak (z) | Brain region | Direction |
|---|---|---|---|---|---|---|
| 1 | 68840 | 13 | -71 | -8 | Right Lingual Gyrus | down |
| 2 | 2184 | 19 | -67 | 40 | Right Precuneus | down |
| 3 | 1712 | -17 | -33 | 4 | Left Thalamus **AND** Pulvinar | down |
| 4 | 1344 | -41 | -67 | -32 | Left Pyramis | up |
| 5 | 1064 | 47 | 5 | -22 | Right Superior Temporal Gyrus **AND** BA 38, Middle Temporal Gyrus | down |
| 6 | 976 | 29 | -7 | 8 | Right Lentiform Nucleus **AND** Putamen | up |
| 7 | 864 | 35 | 43 | 22 | Right Middle Frontal Gyrus **AND** BA 10 | up |
| 8 | 864 | -45 | -3 | -10 | Left Superior Temporal Gyrus | down |
| 9 | 760 | -1 | -31 | 26 | Left Cingulate Gyrus **AND** BA 23 | up |
| 10 | 720 | 9 | -13 | 38 | Right Cingulate Gyrus | down |
| 11 | 672 | -21 | -7 | -10 | Left Parahippocampal Gyrus **AND** Amygdala | down |
| 12 | 624 | -63 | -17 | 30 | Left Postcentral Gyrus **AND** BA 3 | down |
| 13 | 616 | 7 | -27 | 0 | Right Thalamus | down |
| 14 | 480 | 49 | -47 | 46 | Right Inferior Parietal Lobule **AND** BA 40 | up |
| 15 | 472 | 31 | 17 | -12 | Right Inferior Frontal Gyrus **AND** BA 47 | down |
| 16 | 456 | 3 | 17 | 0 | Right Caudate | down |
| 17 | 440 | -35 | -15 | 48 | Left Precentral Gyrus **AND** BA 4 | down |
| 18 | 432 | -45 | -63 | 44 | Left Inferior Parietal Lobule | up |
| 19 | 408 | -27 | 21 | -8 | Left Inferior Frontal Gyrus **AND** BA 47 | down |
| 20 | 400 | 15 | -1 | 0 | Right Lentiform Nucleus **AND** Lateral Globus Pallidus | up |
| 21 | 368 | -31 | -13 | 4 | Left Lentiform Nucleus **AND** Putamen | up |
| 22 | 328 | -5 | -41 | -32 | Left Nodule | up |
| 23 | 320 | -29 | 19 | 8 | Left Insula | up |
| 24 | 304 | -23 | -59 | -24 | Left Culmen | up |
| 25 | 296 | 13 | -71 | -38 | Right Inferior Semi-Lunar Lobule | down |
| 26 | 288 | -13 | -19 | 40 | Left Cingulate Gyrus | down |
| 27 | 240 | -5 | -23 | 58 | Left Medial Frontal Gyrus, BA 6 | up |
| 28 | 224 | -43 | -51 | 34 | Left Supramarginal Gyrus | up |
| 29 | 216 | 37 | 17 | 46 | Right Middle Frontal Gyrus | up |

| | | | | | | |
|---|---|---|---|---|---|---|
| 30 | 216 | -29 | -41 | 50 | Left Inferior Parietal Lobule | down |
| 31 | 208 | -49 | -15 | -4 | Left Superior Temporal Gyrus | down |
| 32 | 208 | -13 | -33 | 40 | Left Cingulate Gyrus | down |
| 33 | 200 | 39 | 7 | -10 | Right Superior Temporal Gyrus **AND** BA 13 | down |
| 34 | 200 | 45 | -43 | 10 | Right Superior Temporal Gyrus | down |
| 35 | 184 | -45 | -37 | 52 | Left Inferior Parietal Lobule | down |
| 36 | 176 | -21 | 49 | 30 | Left Superior Frontal Gyrus **AND** BA 9 | up |
| 37 | 176 | 25 | 5 | 56 | Right Superior Frontal Gyrus **AND** BA 6 | up |
| 38 | 168 | 35 | 35 | 34 | Right Middle Frontal Gyrus **AND** BA 9 | up |
| 39 | 160 | 29 | -57 | -38 | Right Cerebellar Tonsil | up |

**Table 5.** Contrast (post - pre) for the left PA**.** Coordinates are in Talairach-Tournoux space. BA = Brodmann Area. Only clusters of 20 or more adjacent voxels were retained.

| | **Volume ($mm^3$)** | **Peak (X)** | **Peak (y)** | **Peak (z)** | **Brain region** | **Direction** |
|---|---|---|---|---|---|---|
| 1 | 28184 | 43 | -57 | 6 | Right Middle Temporal Gyrus | down |
| 2 | 3968 | 53 | 1 | 40 | Right Middle Frontal Gyrus | down |
| 3 | 2688 | -45 | -33 | 26 | Left Inferior Parietal Lobule | down |
| 4 | 2424 | -35 | -23 | -12 | Left Parahippocampal Gyrus | down |
| 5 | 1824 | -19 | -9 | 66 | Left Superior Frontal Gyrus **AND** BA 6 | down |
| 6 | 1784 | 63 | -17 | 26 | Right Postcentral Gyrus | down |
| 7 | 1632 | 29 | -59 | 46 | Right Superior Parietal Lobule | down |
| 8 | 1048 | 19 | 55 | 32 | Right Superior Frontal Gyrus **AND** BA 9 | up |
| 9 | 960 | 25 | -71 | -4 | Right Lingual Gyrus **AND** BA 19 | down |
| 10 | 808 | 41 | -49 | -16 | Right Fusiform Gyrus **AND** BA 37 | down |
| 11 | 728 | 41 | -33 | 58 | Right Postcentral Gyrus | down |
| 12 | 712 | 21 | -15 | 60 | Right Middle Frontal Gyrus | down |
| 13 | 480 | -11 | -79 | -6 | Left Lingual Gyrus **AND** BA 18 | down |
| 14 | 448 | -13 | -29 | -10 | Left Culmen | up |
| 15 | 440 | -33 | -7 | 58 | Left Middle Frontal Gyrus | down |
| 16 | 408 | 45 | -5 | -8 | Right Superior Temporal Gyrus | down |
| 17 | 400 | 17 | -21 | 16 | Right Thalamus **AND** Lateral Posterior Nucleus | down |
| 18 | 368 | -5 | -19 | 20 | Left Thalamus | up |
| 19 | 352 | 23 | -57 | 12 | Right Posterior Cingulate | down |
| 20 | 312 | 7 | -5 | 2 | Right Thalamus **AND** Ventral Anterior Nucleus | down |
| 21 | 312 | 11 | -67 | -2 | Right Lingual Gyrus **AND** BA 18 | up |
| 22 | 280 | -25 | -47 | -2 | Left Parahippocampal Gyrus **AND** BA 19 | down |
| 23 | 256 | 11 | -35 | 8 | Right Thalamus | up |
| 24 | 256 | -45 | -29 | 2 | Left Superior Temporal Gyrus | down |
| 25 | 248 | 43 | 27 | -12 | Right Inferior Frontal Gyrus | up |
| 26 | 248 | -9 | -19 | 4 | Left Thalamus **AND** Mammillary Body | up |
| 27 | 240 | 33 | 47 | -6 | Right Middle Frontal Gyrus | up |
| 28 | 240 | -17 | 9 | 2 | Left Lentiform Nucleus **AND** Putamen | up |
| 29 | 240 | -1 | 49 | 38 | Left Medial Frontal Gyrus, BA 8 | up |
| 30 | 240 | 11 | -41 | 36 | Right Cingulate Gyrus **AND** BA 31 | down |
| 31 | 216 | 23 | -79 | -22 | Right Declive | down |
| 32 | 216 | 17 | -73 | 36 | Right Precuneus | down |

| | | | | | | |
|---|---|---|---|---|---|---|
| 33 | 208 | 57 | -5 | 0 | Right Superior Temporal Gyrus **AND** BA 22 | down |
| 34 | 200 | -27 | -5 | -16 | Left Parahippocampal Gyrus **AND** Amygdala | down |
| 35 | 200 | 17 | 39 | 42 | Right Superior Frontal Gyrus | down |
| 36 | 192 | 53 | 31 | 20 | Right Middle Frontal Gyrus | down |
| 37 | 184 | 5 | -37 | -52 | - | down |
| 38 | 184 | 7 | -5 | -8 | Right Lentiform Nucleus | down |
| 39 | 176 | -29 | -71 | -8 | Left Lingual Gyrus **AND** BA 18 | down |
| 40 | 168 | 27 | -53 | -20 | Right Culmen | down |
| 41 | 168 | 21 | 3 | -14 | Right Subcallosal Gyrus **AND** BA 34 | down |
| 42 | 160 | 13 | -23 | -14 | Right Culmen | up |
| 43 | 160 | 49 | -21 | 2 | Right Superior Temporal Gyrus | down |
